# PanSVmerger: a flexible pipeline for merging multiallelic structural variants in pangenome graphs

**DOI:** 10.64898/2026.08.13.744739

**Authors:** Ting Yang, Junzhe Shi, Quanyu Chen, Dongya Wu, Xinjiang Tan, Jue Ruan, Chentao Yang

## Abstract

**Summary:** Pangenome graphs capture extensive genetic diversity but introduce analytical challenges due to the redundant representation of structural variations (SVs). While existing tools effectively address cross-sample redundancy or cross-locus redundancy, none specifically target the intra-locus allelic redundancy inherent to pangenome graphs. Here, we present PanSVmerger, an open-source tool designed to consolidate redundant multiallelic SVs within individual loci using three complementary clustering strategies: adaptive k-mer-based Jaccard distance, global alignment distance via VSEARCH, and length distribution. Validation on HPRC pangenome data demonstrates that PanSVmerger effectively reduces multiallelic complexity (e.g., AC ≥ 3 loci from 62.4% to 4.7% using Strategy A) with a modest trade-off: Recall decreased from 97.13% to 93.58%, while precision improved from 94.95% to 96.56%, yielding an overall F1-score of 95.05%. These results demonstrate that PanSVmerger effectively consolidates redundant allele representations with only a minimal loss of sensitivity, making it well-suited for downstream applications that require clean, non-redundant variants.

**Availability and implementation:** PanSVmerger is implemented in Python 3.8+ and freely available under the MIT license at GitHub: https://github.com/tingting100/PanSVmerger. The software requires vcflib, bcftools, and optionally VSEARCH. Comprehensive documentation and tutorials are provided.

## 1 Introduction

Recent advances in pangenome initiatives, including the Human Pangenome Reference Consortium (HPRC) (Liao *et al*., 2023) and Chinese Pangenome Consortium (CPC) (Gao *et al*., 2023), Arab Pangenome Reference (UPR) (Nassir *et al*., 2025) have driven the adoption of graph-based pangenomes constructed using tools like Minigraph (Li *et al*., 2020), Minigraph-Cactus (Hickey *et al*., 2024) and PanGenome Graph Builder (PGGB) (Garrison *et al*., 2024). These graphs capture extensive structural variation across diverse populations, significantly advancing variant discovery beyond traditional linear reference genomes. However, a major challenge persists: the representation of structural variations (SVs) in pangenome graphs frequently results in highly complex multiallelic loci where a single underlying SV event is represented as multiple redundant allele records, increasing computational burden and complicating downstream analyses, including genome-wide association studies (GWAS), functional annotation, and population genetics (Liao *et al*., 2023; Nie *et al*., 2026). For instance, even with base-level aligment, Minigraph still exhibites breakpoint precision issues (Leonard *et al*., 2023). Conversely, reference-free aligners such as PGGB and Cactus achieve base-level resolution but inevitably produce nested variation and substantially higher computational costs (Crysnanto *et al*., 2022). Together, these technical factors create representational redundancy that inflate multiallelic loci, obscuring true allele frequencies and complicating biological interpretation.

Recognizing these challenges, the genomics community has developed several tools to address SV redundancy, though they primarily focus on other dimensions of the problem (**Table 1**). Cross-sample merging tools, such as SURVIVOR (Jeffares *et al*., 2017) and bcftools (Danecek *et al*., 2021) consolidate variants across different individuals into a unified cohort-level VCF. Cross-locus utilities, like Truvari collapse (English *et al*., 2022) compare variants at disparate genomic positions to determine if they stem from the same biological event. More recently, PanPop (Zheng *et al*., 2024) was introduced to merge SVs across multiple callers or individuals. In contrast, the vcfwave (Garrison *et al*., 2022) offers a decomposition strategy, it splitting complex, nested ALT alleles into simpler “primitive” variants via pairwise alignment with BiWFA (Marco-Sola *et al*., 2023). While this normalization improves variant representation for certain applications like breakpoint annotation, it fundamentally fragments the original relationship between alleles, rendering the population allele frequency (AF) of the original structural variant undefinable. To our knowledge, no existing tool addresses the fundamental challenge of consolidating redundant ALT alleles within a single multiallelic VCF record while preserving the true population allele frequency—a problem that is particularly exacerbated by the nested and fragmented variant representations produced by pangenome graph.

**Table 1.** Comparison of SV merging and normalization tools.

| Tool | Primary purpose | Algorithm basis | Output type |
| --- | --- | --- | --- |
| SURVIVOR | Multi-caller & cross-sample merging | Position overlap | Multiallelic VCF |
| bcftools merge | cross-sample merging | Exact position & allele match | Multiallelic VCF |
| PanPop | Multi-caller / cross-sample merging | Sequence-aware realignment | Biallelic VCF |
| Truvari collapse | Cross-locus SV collapsing | Sequence + Length similarity | Collapsed VCF |
| vcfwave | Complex allele decomposition | BiWFA pairwise alignment | Decomposed VCF |
| PanSVmerger (this work) | Intra-locus allelic redundancy consolidation | Sequence similarity + Length | Non-redundant VCF |

To bridge this critical methodological gap, we developed PanSVmerger, the first dedicated pipeline designed to resolve redundant multiallelic SV representations in pangenome graphs. PanSVmerger provides three complementary clustering strategies that leverage sequence composition and length distribution at the allele level. This consolidation enables vital downstream applications: the resulting resolved SV genotypes can be directly integrated into standard GWAS tools, phylogenetic methods, and population genetics software that require simple, robust allele counts.

## 2 Methods

### 2.1 Overview of PanSVmerger pipeline

PanSVmerger accepts multi-sample VCF files containing SVs from pangenome graphs (e.g., Minigraph-Cactus output) as input. The workflow comprises three main stages:

#### Stage 1

Regional filtering. PanSVmerger optionally excludes SVs located in complex genomic regions, including centromeres, telomeres using user-provided BED files.

#### Stage 2

SV locus extraction. The pipeline identifies qualifying multi-allelic loci where the maximum length difference between the reference and any alternate allele sequence is ≥ 50 bp. This threshold aligns with standard SV definitions, focusing the downstream clustering on genuine structural variants. Crucially, once a locus is selected based on this criterion, all associated ALT sequences—including secondary nested or shorter alleles—are extracted to ensure comprehensive intra-locus resolution.

#### Stage 3

Allele clustering. All allele sequences at each qualifying locus are consolidated using one of three complementary strategies: k-mer-based adaptive clustering using Jaccard index (Strategy A), global alignment-based clustering using VSEARCH (Strategy B), or pure length-based clustering (Strategy C).

Following clustering, the VCF file is automatically updated through representative allele selection (choosing the longest sequence per cluster), genotype recoding to cluster-based IDs, and the recalculation of population statistics, including Allele Count (AC), Allele Number (AN), and Allele Frequency (AF) (**Fig. 1**).

**Fig. 1.**
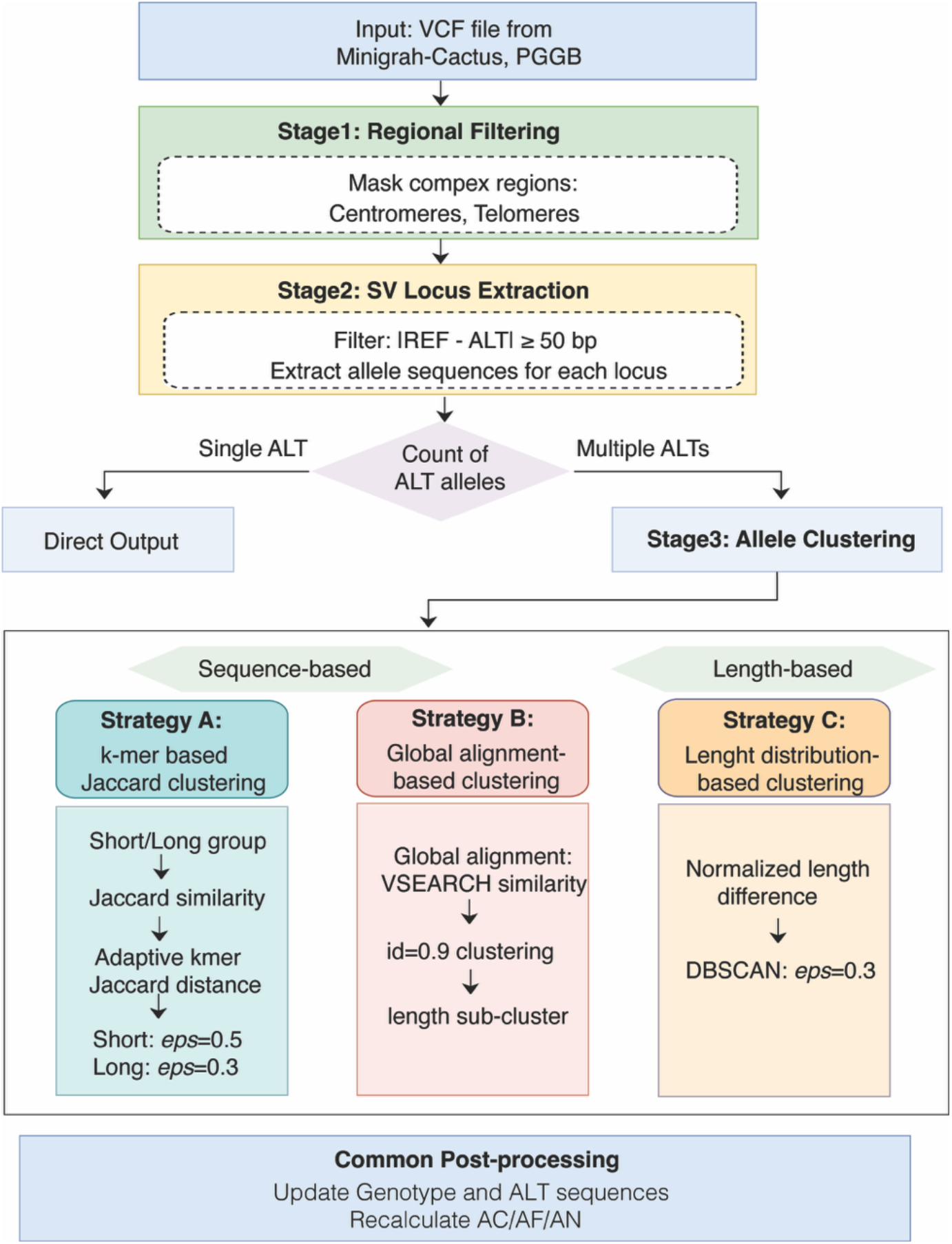
PanSVmerger workflow.

For clustering, PanSVmerger identifies genomic positions where ALT and REF sequences differ by ≥50 bp. At each qualifying locus, all allele sequences are extracted and clustered based on sequence similarity and/or length differences. Following clustering, the VCF is updated through: (a) selecting representative alleles (longest sequence per cluster); (b) updating REF to cluster 0’s representative; (c) recoding genotypes to cluster-based IDs; and (d) recalculating population statistics (AC, AN, AF). This ensures VCF format compatibility for downstream analyses while eliminating redundancy.

#### 2.1.1 Strategy A: K-mer-based adaptive clustering (Jaccard method)

This strategy quantifies sequence similarity using an adaptive k-mer Jaccard distance metric (Bonnici *et al*., 2022). For each sequence pair (seq_i, seq_j), the distance is defined as:

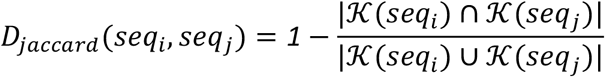

where *K*(*seq*) represents the set of k-mers extracted from the sequence. To accommodate varying sequence lengths, we implemented adaptive k-mer selection: k=1 for ultra-short sequences (≤5 bp), k=2 for short (6-10 bp), k=3 for medium (11-20 bp), and k=5 for long (>20 bp) sequences.

An adaptive weighting parameter α automatically balances k-mer similarity against length differences based on sequence length variability at each locus:

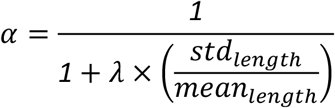

where λ (default: 2.5) is a user-adjustable scaling factor controlling the sensitivity of α to length variation. The term *std*_*length*_/*mean*_*length*_ is the coefficient of variation (CV) of allele lengths at a given locus, which measures length dispersion relative to the mean. We analyzed the distribution of length variation across all loci (median CV = 0.103) and selected λ=2.5 to allow a smooth transition of α from 0.80 (typical loci) to 0.57 (moderately variable loci), ensuring that sequence similarity dominates when length differences are small while length information gains weight as variation increases. The final distance matrix combines both metrics:

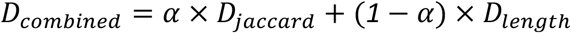

Where:

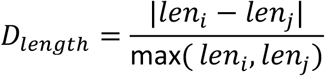

To account for the different clustering characteristics of short and long alleles, sequences are partitioned into short (<50 bp) and long (≥50 bp) groups. For short sequences, DBSCAN clustering (eps=0.5, min_samples=1) is applied; for long sequences, a stricter eps=0.3 is used. These epsilon values were empirically selected based on Jaccard distance distributions observed in preliminary analyses (short: inflection point ∼0.4–0.6; long: stricter threshold to preserve diversity). Noise points are reassigned to unique singleton clusters.

#### 2.1.2 Strategy B: Global alignment-based clustering (Vsearch method)

For base-pair resolution similarity assessment, we employed VSEARCH (v2.22.1) (Rognes *et al*., 2016) with the --cluster_fast algorithm, which performs global pairwise alignment: *vsearch --cluster_fast {input} --id 0*.*9 --strand both --uc {output}*.*uc -- threads {threads}*. By default, sequences are clustered at a 90% identity threshold. To handle nested SVs, we implemented hierarchical refinement within each VSEARCH cluster: sequences that differ in length by more than 10% from the longest sequence are partitioned to distinct sub-clusters. This two-tier approach ensures that sequences sharing high global similarity but possessing substantial length differences (e.g., nested SVs) are appropriately separated, thereby preserving biologically distinct alleles. For example, a 500 bp deletion nested within a 1000 bp deletion would be clustered together by global alignment due to high sequence identity over the aligned region but will be correctly separated by the 10% length difference threshold, preserving biologically distinct alleles.

#### 2.1.3 Strategy C: Length distribution-based clustering (Length method)

As a computationally rapid alternative for large datasets, we implemented a pure length-based clustering strategy based using on normalized length difference:

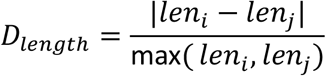

This generates a distance matrix bound between 0 and 1, which serves as the input for DBSCAN clustering (eps=0.3, min_samples=2). Length-based clustering serves as an efficient screening tool for massive datasets where processing speed is prioritized, and it is highly effective for SV classes where length is the primary distinguishing feature.

### 2.2 Data sources

To evaluate PanSVmerger on realistic pangenome topologies, we utilized two complementary pangenome graph datasets. First, we reconstructed the HPRC Minigraph-Cactus pangenome graph using T2T-CHM13 (Nurk *et al*., 2022) as the linear reference genome, incorporating assemblies from HG002 (Hansen *et al*., 2025), HG005, and GRCh38 to yield a total of 95 haplotypes. The variants from HG002 were specifically extracted for downstream benchmarking. Second, we utilized the publicly available HPRC PGGB pangenome graph downloaded from the official human pangenomics repository (https://github.com/human-pangenomics/hpp_pangenome_resources). For performance assessment, precision-recall metrics for HG002 were generated by comparing our consolidated variant calls against the CHM13v2.0_HG2-T2TQ100-V1.1 SV benchmark set, treating the NIST draft benchmark files as the ground truth (https://ftp-trace.ncbi.nlm.nih.gov/ReferenceSamples/giab/data/AshkenazimTrio/analysis/NIST_HG002_DraftBenchmark_defrabbV0.020-20250117/).

### 2.3 Implementation details

PanSVmerger is implemented in Python 3.8+ with modular architecture. All three clustering strategies share unified VCF input/output via pysam, genotype recoding, and statistical recalculation. High-performance parallel processing is supported via concurrent.futures module for multi-threaded execution of independent loci across a configurable thread count (default: 4). The pipeline includes command-line interfaces for each strategy, allowing full customizability of key parameters (*eps, λ, k*, identity threshold).

Special handling ensures: (i) The REF sequence is strictly pinned to cluster 0; (ii) Noise points from DBSCAN are retained as unique singleton clusters rather than being erroneously discarded; (iii) Short sequences (<50 bp) within a multiallelic locus receive adjusted distance thresholds; (iv) Sequences shorter than the designated k-mer size fall back to k=1; and (v) VCF metadata and INFO fields (AC, AN, AF) are accurately recalculated to maintain downstream compatibility.

## 3 Results

### 3.1 Performance on HPRC pangenome data

We evaluated PanSVmerger on SV sets from the HPRC pangenome, which comprises 47 diverse human genomes called with Minigraph-Cactus and PGGB. We examined the distribution of allele counts (AC) per locus in the raw pangenome graphs (**Fig. 2a**). For the Minigraph-Cactus graph, 49,944 loci (62.85%) harbored three or more alternate alleles, while for the PGGB graph, 64,226 loci (65.05%) showed AC ≥3. The prevalence of multiallelic loci with high allele numbers indicates widespread representational redundancy during graph construction—a consequence of breakpoint imprecision, nested variation, and other technical factors that can dilute the observed allele frequency of the underlying SV event and complicate downstream analyses.

**Fig. 2.**
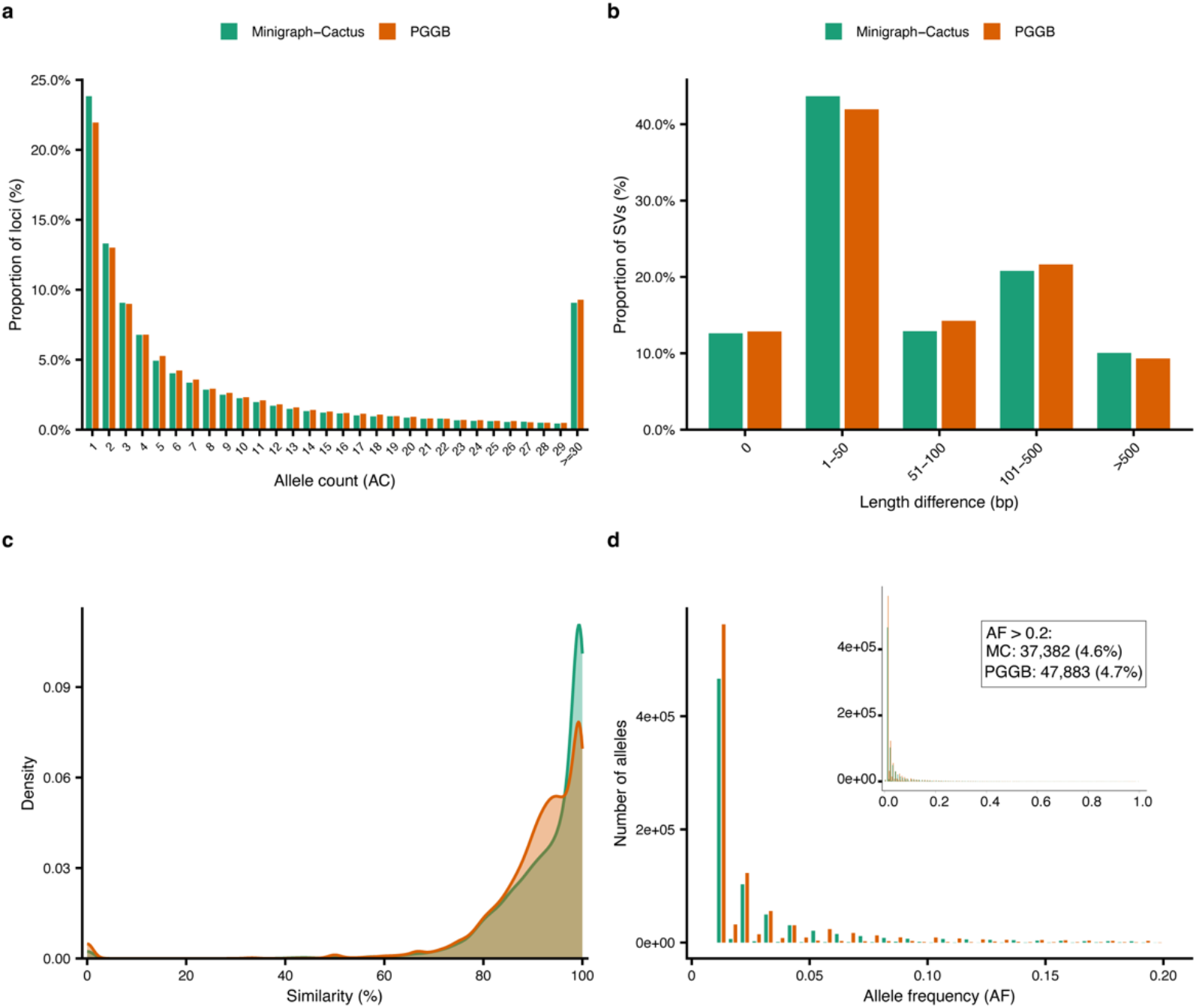
Characteristics of multiallelic redundancy in raw pangenome graphs. (a) Distribution of allele counts (AC) in the original multi-sample VCF from Minigraph-Cactus, prior to merging. The x-axis shows allele count categories, and the y-axis reports the number of variants per category, revealing the prevalence of multi-allelic records in the dataset. (b) Distribution of length differences between each alternate allele and the longest allele at the same locus, with x-axis categories (0, 1–50, 51–100, 101–500, >500 bp) and y-axis showing the number of alleles per category. This highlights the proportion of alleles with substantial length variation that would benefit from length-based clustering. (c) Density distribution of sequence similarity between each alternate allele and the longest allele at the same locus, revealing that a large fraction of alleles exhibits high sequence similarity. (d) Allele frequency (AF) distribution of alternate alleles in raw graphs.

We next evaluated the distribution of length differences between each alternate allele and the reference allele at the same locus (**Fig. 2b**). In the Minigraph-Cactus graph, 12.6% of alleles showed zero length difference relative to the reference sequence, while the largest fraction (43.7%) fell into the 1–50 bp range; collectively, 56.3% of all alternate alleles differed by ≤50 bp. Alleles with length differences of 51–100 bp, 101– 500 bp, and >500 bp accounted for 12.9%, 20.8%, and 10.1%, respectively. A nearly identical length distribution was observed for the PGGB graph. These findings suggest that the majority of graph-derived alleles exhibit minor length differences (≤50 bp) that likely represent technical redundancy rather than distinct biological events, making them ideal candidates for consolidation.

To confirm this hypothesis, we assessed sequence similarity among all alternate alleles within each locus using an all-vs-all pairwise global alignment (**Fig. 2c**). In the Minigraph-Cactus graph, 69.9% of allele pairs shared >90% sequence identity, with the PGGB graph displaying a similar proportion (65.2%). This high prevalence of near-identical sequences, coupled with the minimal length variations, suggests that many of these alleles may represent fragmented records of the same underlying genomic event. This interpretation is further supported by the extreme skew in allele frequencies (**Fig. 2d**), where 57.9% of alleles in the Minigraph-Cactus graph and 55.3% in the PGGB graph were confined to the lowest frequency bin (AF = 0.0125). Across both graphs, over 80% of alleles exhibited an AF < 0.05 (PGGB: 81.7%; Minigraph-Cactus: 80.4%), consistent with the view that the same SV events are often fragmented into multiple low-frequency records, diluting the true allele frequency of the underlying variant. This fragmentation likely stems from technical factors such as breakpoint imprecision and graph traversal ambiguities, rather than reflecting genuine allelic diversity. These findings highlight the importance of consolidating redundant allele representations to recover accurate allele frequencies for downstream analyses.

To demonstrate the efficacy of PanSVmerger in mitigating this redundancy, we compared the intra-locus AC distributions before and after consolidation (**Fig. 3**). Following application to the Minigraph-Cactus graph, the proportion of highly complex loci (AC ≥ 3) dropped drastically from 62.4% to 4.7% using Strategy A (Jaccard method), 26.3% using Strategy B (VSEARCH method), and 11.3% using Strategy C (Length method). Concurrently, the proportion of clean biallelic loci increased from a baseline of 24.1% to 80.2%, 62.1%, and 72.9%, respectively. Strategy A achieved the most aggressive consolidation, reducing the average AC from 9.7 to 1.3, while Strategy B was the most conservative (average AC: 2.6), and Strategy C delivered an intermediate balance (average AC: 1.7). Consistent consolidation trends were observed for the PGGB graph, where complex loci (AC ≥ 3) were reduced from 64.4% to 6.1%, 29.8%, and 11.2% across the three strategies, respectively. These results underscore the flexibility of PanSVmerger, allowing users to select consolidation thresholds tailored to specifical analytical requirements.

**Fig. 3.**
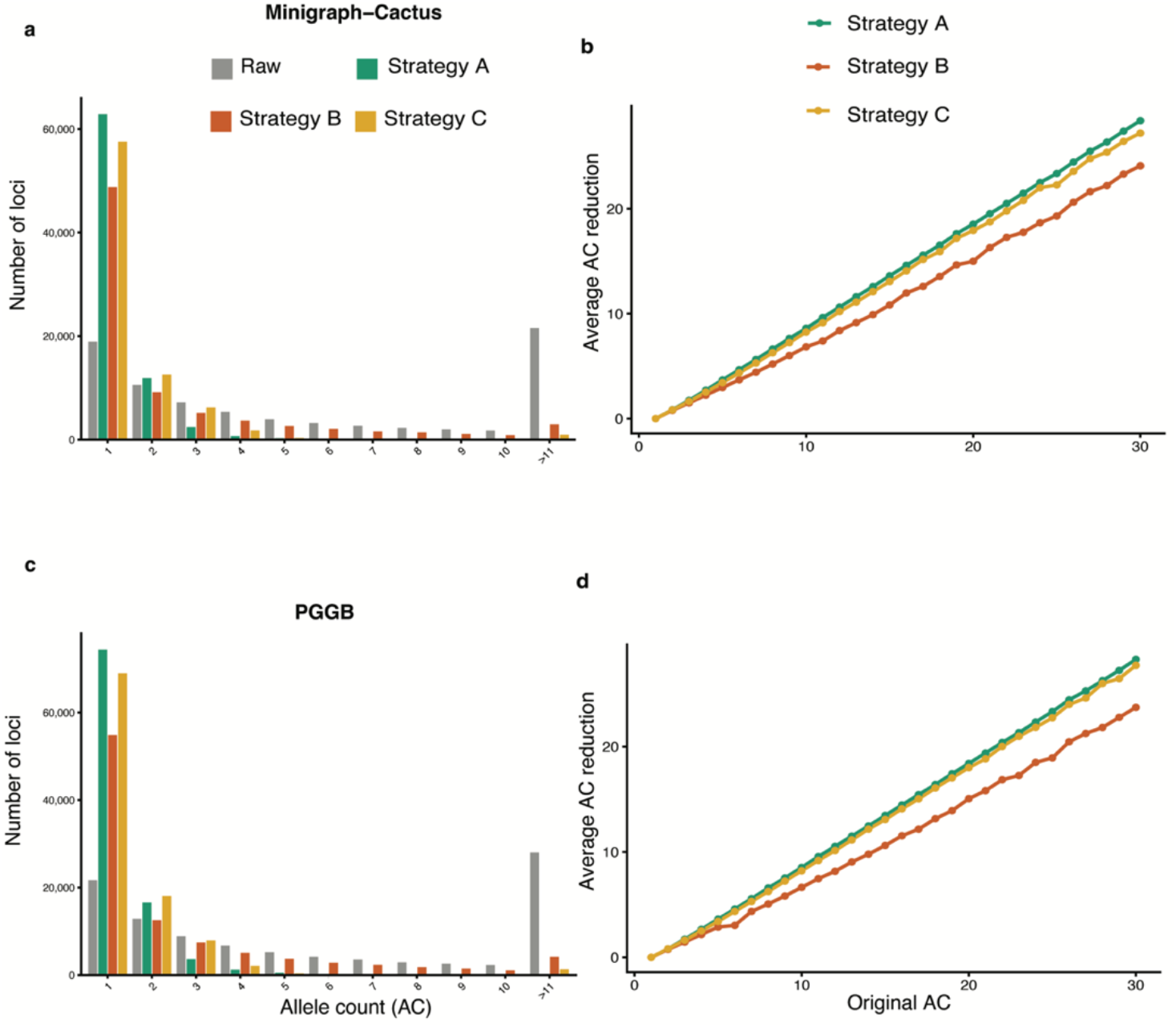
Comparison of AC distributions before and after PanSVmerger merging. (a, c) Bar plots show the number of loci with different allele counts (AC) for the raw graph and for each of the three clustering strategies in Minigraph-Cactus and PGGB, respectively. (b, d) Average AC reduction (raw AC minus merged AC per locus) for each strategy in Minigraph-Cactus and PGGB, respectively. Strategies: A, Jaccard-based; B, VSEARCH-based; C, Length-based.

When stratified by specific SV classes, we found that for deletion, the predominant SV type in our dataset (58.3%), all three methods achieved >90% consolidation efficiency for alleles with <10% length difference, while successfully preserving >95% of truly divergent alleles with >20% length difference. For insertions, which present elevated sequence complexity due to repetitive content, Strategy A significantly outperformed the other methods, consolidating 87.3% of highly similar alleles (>95% identity) while safely preserving 92.1% of highly divergent sequences (<80% identity).

### 3.2 Benchmarking on HG002 truth set

To evaluate genotyping accuracy, we benchmarked the three PanSVmerger strategies against the Genome in a Bottle (GIAB) HG002 high-confidence truth set using Truvari bench with parameters -r 100 -p 0.8 -O 0 -P 0.8 (the PGGB graph does not include HG002 samples and was omitted from this validation). Raw Minigraph-Cactus SV calls achieved 94.95% precision and 97.13% recall (F1 = 96.03%), serving as the baseline against which all strategies were compared. Both sequence-based strategies, Strategy A (Jaccard-based) and Strategy B (VSEARCH-based), exhibited highly comparable overall accuracy (F1: 95.05% vs. 95.20%) and substantially outperformed the raw graph calls in precision (96.56% and 96.32%, respectively). Strategy C (Length-based), while more conservative, maintained precision at 94.35% with 90.54% recall (F1 = 92.41%) (**Table 2**). These results demonstrate that PanSVmerger effectively consolidates intra-locus allelic redundancy while preserving genuine variant signals. The complementary strengths of the two sequence-based strategies offer users flexibility in balancing precision and recall according to specific analytical priorities.

**Table 2.**
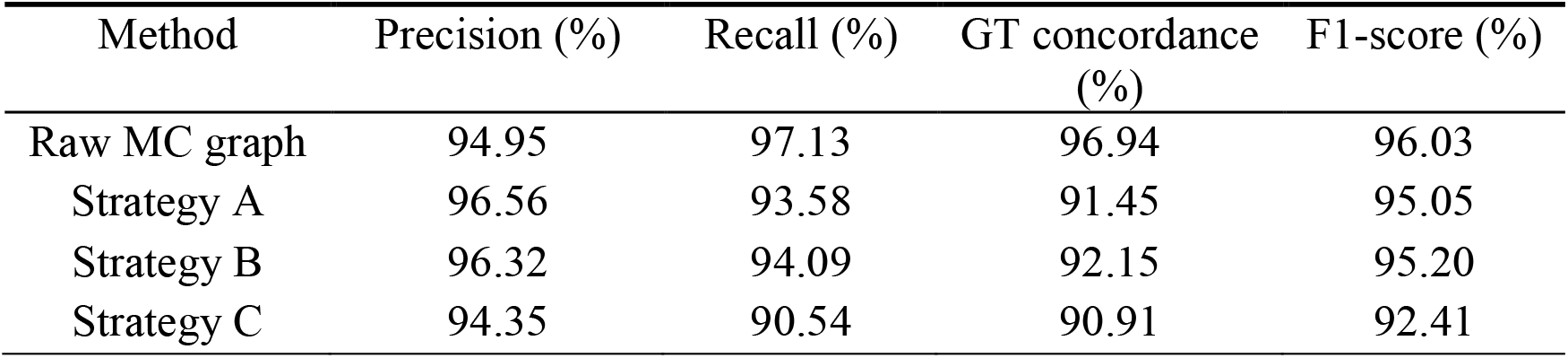
Genotyping performance metrics relative to the GIAB HG002 truth set. All values are reported as percentages calculated against raw Minigraph-Cactus SV calls.

To further benchmark PanSVmerger against existing SV merging tools, we compared its performance with three widely used approaches: PanPop, Truvari collapse, and vcfwave. For PanPop and Truvari, we first normalized the multiallelic sites into biallelic variants using bcftools norm -m -any. In contrast, vcfwave was applied directly to the raw Minigraph-Cactus multiallelic VCF with the –L 10000 option to decompose complex alleles into primitive variants, followed by bcftools and vcfcreatemulti. All tools were evaluated on the same HG002 benchmark set using identical Truvari bench parameters to ensure a fair comparison (**Fig. 4)**.

**Fig. 4.**
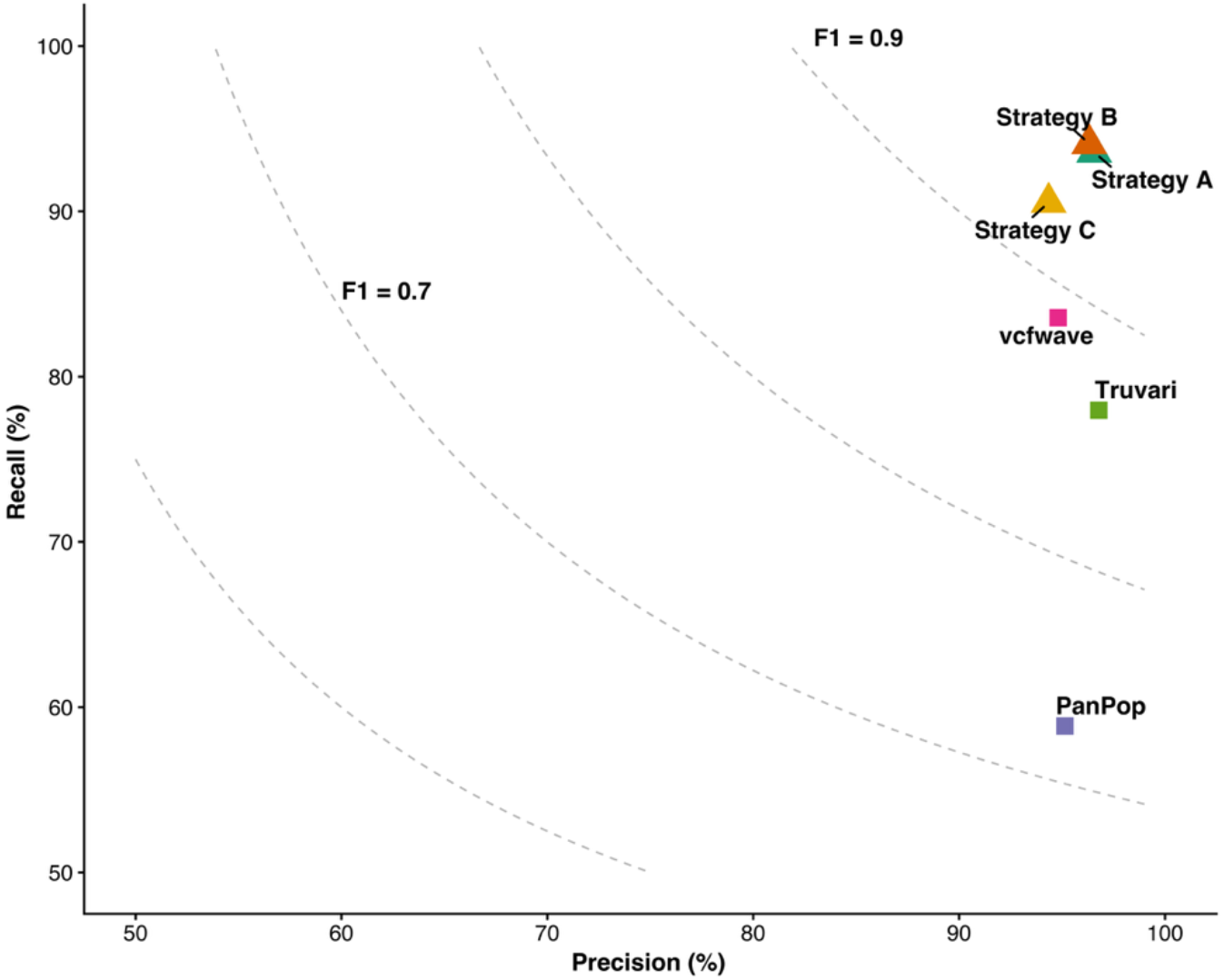
Precision-recall comparison of SV merging tools on the HG002 benchmark set. Each point represents a tool. Gray dashed curves indicate F-measure contours at 0.6–0.9 intervals. PanSVmerger (Strategy A, B and C) are shown as large colored points. Raw graph calls (gray hollow circle) are shown as baseline. PanPop, vcfwave, and Truvari are included for comparison.

Among the existing tools, Truvari achieved the highest precision (96.78%) but at the cost of substantially lower recall (77.97%), resulting in an F1 of 86.36%. Vcfwave attained moderate performance with 94.81% precision and 83.57% recall (F1 = 88.84%), reflecting its decomposition-based approach that fragments complex alleles into primitive records. PanPop exhibited the lowest recall (58.86%) and F1 (72.72%), despite moderate precision (95.14%). In contrast, PanSVmerger (Strategy B) achieved the highest F1 (95.20%) among all tools, outperforming Truvari by 8.84 percentage points, vcfwave by 6.36 percentage points, and PanPop by 22.48 percentage points.

These results demonstrate that PanSVmerger provides a more balanced and effective approach for consolidating intra-locus allelic redundancy than existing methods, achieving superior overall accuracy without sacrificing recall.

### 3.3 Case Study: Resolving Allelic Redundancy at the LPA Locus

To evaluate PanSVmerger’s performance in complex genomic regions, we conducted a targeted case study of the Lipoprotein A (*LPA*) gene locus (chr6:161.78– 162.01 Mb, T2T-CHM13 v2.0) (Volgman *et al*., 2024). This region is characterized by highly polymorphic Kringle IV type 2 (KIV-2) tandem repeats, which frequently cause severe allele fragmentation in graph-based pangenomes due to their intense sequence homology. Within the raw HPRC v1.1 (Minigraph-Cactus) pangenome VCF, we identified a highly fragmented multiallelic site containing 13 alternative alleles (ALTs), which likely represented redundant records of a limited number of underlying allelic stats rather than distinct biological events.

Application of PanSVmerger suceessfully collapsed this complex locus across all three strategies. Strategy A and C prioritized maximum noise reduction, collapsing all 13 redundant ALTs into a single high-confidence representative allele (an 87.5% reduction in alternate allele count), whereas Strategy B adopted a more conservative threshold, merging them into 4 distinct allelic classes (50.0% reduction) to preserve fine-grained variations (**Fig. 5**). The biological validity of this consolidation was supported by an average intra-cluster Jaccard distance of 0.1458, corresponding to an approximately sequence similarity of 84.5% among the merged alleles. This high similarity confirms that the collapsed sequences represent same underlying structural events across the KIV-2 motifs. By converting fragmented, noisy loci into structured multiallelic sites, PanSVmerger effectively aggregates sparse genotypes into high-confidence alleles, boosting effective allele frequency and enhancing the statistical power for downstream cardiovascular association studies.

**Fig. 5.**
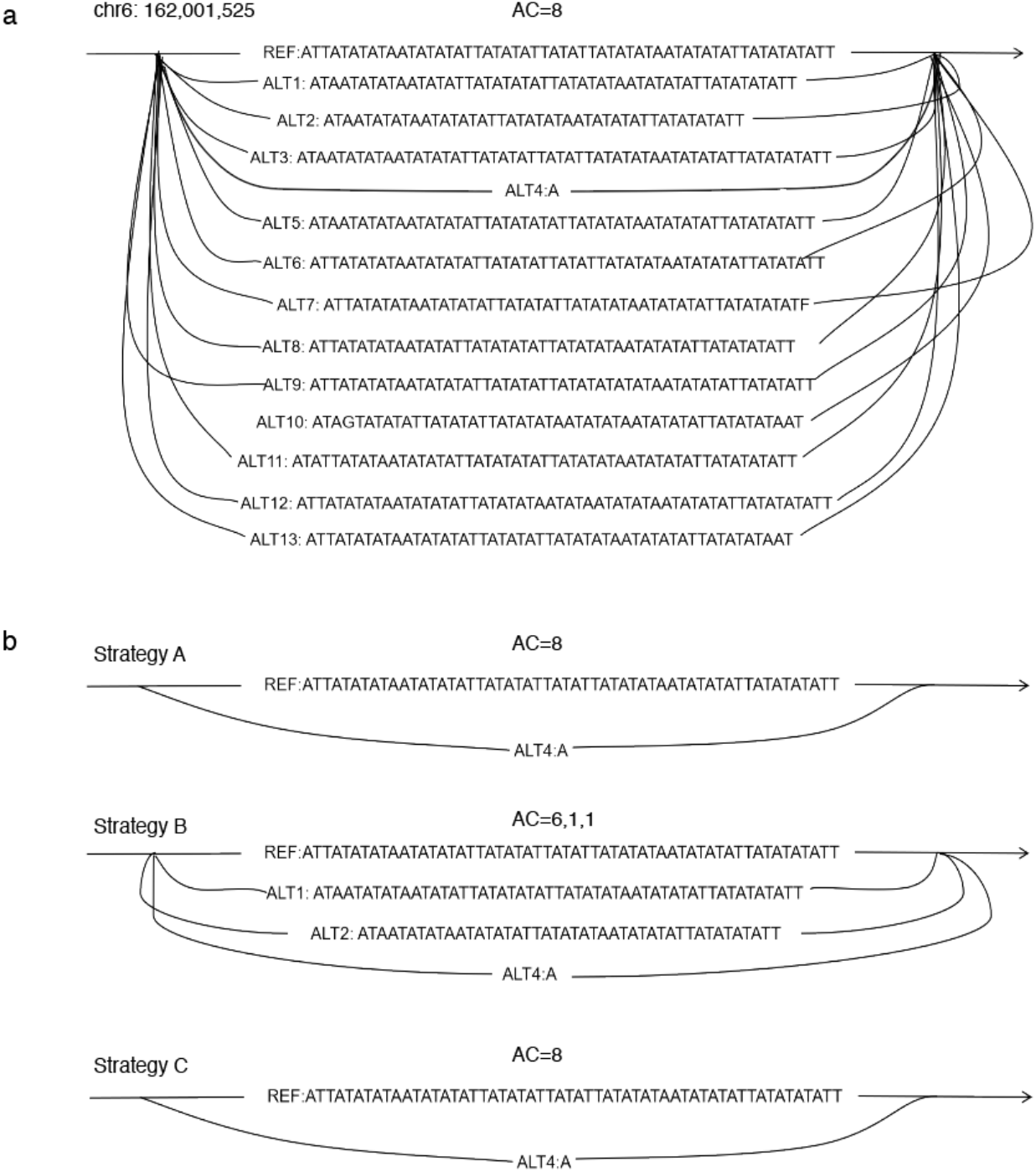
Consolidation of allelic redundancy at the *LPA* gene locus (chr6:161.78–162.01 Mb). (a) Allele distribution in the original Minigraph-Cactus (MC) graph, displaying 13 fragmented alternate alleles. (b) Post-consolidation allele distributions after applying the three PanSVmerger strategies (A: Jaccard, B: Vsearch, C: Length-based). Strategy A and C aggressively collapsed the 8 alleles into a single representative (87.5% reduction), while Strategy B merged them into 4 distinct classes (50.0% reduction)

### 3.4 Computational resource usage

We benchmarked the computational efficiency of the three strategies on chromosome 1 (**Table 3**). Strategy C (Length-based) completed in 14 seconds, reflecting the low computational cost of length-based distance calculation. Strategy A (Jaccard) required approximately 20 minutes, with a CPU usage of 94.73%, indicating efficient utilization of the allocated cores. Strategy B (VSEARCH) was the most computationally intensive, taking 3.5 hours and achieving 242.72% CPU usage, consistent with the overhead of pairwise global alignment. Peak memory usage remained moderate across all strategies (0.50–0.60 GB), making PanSVmerger suitable for resource-constrained environments. These results highlight the trade-off between merging accuracy and computational cost, allowing users to select the most appropriate strategy based on their specific requirements.

**Table 3.** Computational resource usage of PanSVmerger strategies.

| Method | CPU<br>cores | CPU time<br>(min) | CPU<br>usage (%) | Peak memory<br>(GB) |
| --- | --- | --- | --- | --- |
| Strategy A | 3 | 18.88 | 94.73 | 0.50 |
| Strategy B | 3 | 516.92 | 242.72 | 0.60 |
| Strategy C | 3 | 0.15 | 65.22 | NA |

## 4 Conclusions and Discussions

Pangenome graphs have emerged as a powerful framework for capturing genetic diversity beyond the constraints of traditional linear reference genomes. For instance, the HPRC draft pangenome reference, constructed via Minigraph-Cactus, has enabled the identification of 29.6 million variants, decreasing small variant discovery errors by 34% and increasing structural variants detected per haplotype by 104% compared to GRCh38-based workflows (Liao *et al*., 2023). Beyond human genomics, the accessibility and scalability of pangenome graph construction tools have accelerated their adoption across a growing number of plant and animal species, including rice (Guo *et al*., 2025), soybean (Yano *et al*., 2025), cattle (Leonard *et al*., 2023), and pigs (Li *et al*., 2024). The Minigraph-Cactus pipeline, in particular, has emerged as a widely adopted method due to its balance between graph complexity and variant discovery performance, supporting multiple genomes and diverse species. However, different graph construction algorithms generate variant callsets with distinct structural properties: PGGB and Cactus achieve base-level resolution but inevitably resolve bubbles with multiple paths (Crysnanto *et al*., 2022), whereas Minigraph-Cactus relies on post-processing tools such as vcfbub and truvari collapse to filter large bubbles and consolidate redundant SVs into a non-redundant callset (He Y, 2025). It is important to note that the alternative alleles observed in pangenome graphs are not biologically false; rather, they represent genuine sequence variation present in individual samples. However, many of these alleles are highly similar in sequence, differ only marginally in length, and occur at extremely low allele frequencies (Fig. 2). In the context of downstream analyses such as GWAS, these redundant representations collectively dilute the allele frequency of the underlying dominant allele, making it difficult to detect genuine genotype–phenotype associations. Consolidating these alleles is therefore not about removing ‘false’ variants, but about recovering the dominant allelic state at each locus to enable accurate allele frequency estimation and robust statistical testing.

PanSVmerger addresses these redundant representations by successfully integrating sequence composition and length metrics to capture complementary dimensions of SV similarity. The utility of PanSVmerger is underpinned by several key design features. Its modular architecture is a significant advantage, allowing users to select the optimal clustering strategy based on their specific research goals, whether they prioritize precision or recall. The tool incorporates an adaptive α parameter that dynamically adjusts to local sequence variance, which substantially reduces the need for intensive manual parameter optimization. Furthermore, the pipeline ensures statistical fidelity by strictly preserving population genetics metrics, such as AC, AN, and AF, which are essential for downstream population studies. Coupled with its seamless compatibility with standard VCF toolchains and high computational scalability—where the length-based method can process large multi-sample cohorts within hours—PanSVmerger serves as a versatile solution for modern pangenomic data analysis.

Despite these strengths, it is important to acknowledge the limitations of the current implementation. First, the current implementation focuses exclusively on intra-locus merging and does not perform cross-locus consolidation. This design choice was made deliberately: Minigraph-Cactus graphs, which are the primary target of PanSVmerger, employ graph pruning and coordinate flattening strategies that effectively anchor SVs to unified genomic intervals (Hickey *et al*., 2024), making cross-locus fragmentation less frequent than in PGGB or raw Minigraph outputs. Second, although the adaptive α parameter reduces manual intervention, optimal performance still depends on user-specified hyper-parameters, such as DBSCAN *eps* or VSEARCH identity thresholds. Third, The optional regional filtering module relies on high-quality genomic annotations (e.g., centromere/telomere coordinates), which may be unavailable for non-model organisms.

Looking ahead, PanSVmerger holds substantial potential for improving SV analysis beyond human genomics, particularly in non-model organisms and complex plant genomes characterized by high repeat content. The non-redundant, high-fidelity SV catalogs generated by our tool will directly improve the reliability of downstream applications, including structural variant-based genome-wide association studies (SV-GWAS), phylogenetic inference, and functional annotation. While our current validation was performed on HPRC population data, which was not specifically designed for GWAS, we envision that applying PanSVmerger to carefully designed population cohorts will enable systematic evaluation of its impact on association studies. Importantly, PanSVmerger outputs a non-redundant VCF that can be directly used for SV genotyping in short-read populations using tools such as PanGenie (Ebler *et al*., 2022) or vg (Garrison *et al*., 2018) without requiring additional decomposition or normalization steps. Future development will focus on incorporating specialized distance metrics for complex copy number variations (CNVs) and variable number tandem repeats (VNTRs). As pangenome graphs rapidly scale across diverse species, tools like PanSVmerger will be indispensable for transforming raw graph outputs into clean, analysis-ready variant catalogs.

## Declarations

### Ethics approval and consent to participate

Not applicable.

### Consent for publication

Not applicable.

### Authors’ contributions

Conceptualization: C.Y., T.Y.; Supervision: C.Y., J.R.; Software: T.Y., J.S.; Funding Acquisition: C.Y., J.R.; Writing—original draft: T.Y., C.Y.; Writing—review & editing: T.Y., C.Y., J.S., Q. C., D. W., J.R., X. T.

### Availability of data and materials

All genomic datasets used in this study are publicly available. The HPRC Minigraph-Cactus and PGGB pangenome graph was downloaded from **https://github.com/human-pangenomics/hpp_pangenome_resources**. No new datasets were generated or analysed during the current study.

## Acknowledgements

We would like to thank DCS Cloud (https://cloud.stomics.tech/) for providing the computational resources and software support necessary for this study.

## Clinical trial number

Not applicable.

